# Characterization of Bergmann Glia-like Progenitors in the Postnatal Mouse Cerebellum by *in vivo* Electroporation and Spatial Transcriptomics

**DOI:** 10.1101/2025.05.20.655231

**Authors:** Kyoka Suyama, Toma Adachi, Minami Mizuno, Kaiyuan Ji, Eriko Isogai, Ikuko Hasegawa, Kayo Nishitani, Masaki Sone, Satoshi Miyashita, Tomoo Owa, Mikio Hoshino

**Affiliations:** Department of Biochemistry and Cellular Biology, National Center of Neurology and Psychiatry (NCNP), Tokyo 187-8502, Japan; Graduate School of Medical and Dental Sciences, Science Tokyo, Tokyo 152-8550, Japan; Department of Biomolecular Science, Faculty of Science, Toho University, Chiba 274-8510, Japan

**Keywords:** Cerebellum, Astroglial cell, Bergmann glia-like progenitor, *in vivo* electroporation, spatial transcriptomics

## Abstract

In the mammalian cerebellum, three types of astroglial cells—Bergmann glial cells (BGs), inner granule cell layer (IGL) astrocytes, and white matter (WM) astrocytes—arise in postnatal timing from two types of progenitors: Bergmann glia-like progenitors (BGLPs) and astrocyte-like progenitors (AsLPs). In contrast to AsLPs, which are commonly observed in other brain regions, BGLPs have not been well studied. Here we investigate their dynamic changes in number, their differentiation abilities and their gene expression profiles during cerebellar development. BGLPs and AsLPs decrease in number as development progresses from P0, and are almost absent by P10. We developed an electroporation-based method to investigate the progeny cells of BGLPs. We found that BGLPs at P6 differentiate into BGs and IGL astrocytes, but not into WM astrocytes, which is consistent with a previous report. However, BGLPs at P0 were observed to differentiate into not only BGs and IGL astrocytes, but also WM astrocytes, indicating that P0 BGLPs possess wider pluripotency than P6 BGLPs. By conducting spatial transcriptomic analysis of the cerebellum at P0 and P6 with over 5,000 probes (Xenium, 5k), we successfully obtained clusters corresponding to BGLPs at P0 and P6, respectively. Further informatics analyses suggested that P0 BGLPs exhibit more stem cell-like features, while P6 BGLPs show a shift toward BG-like characteristics. This study, which includes transcriptome big data, will contribute to understanding the differentiation of BGs and astrocytes, as well as other types of cells, during postnatal cerebellar development.

## Introduction

Astroglial cell diversity is crucial for proper brain function in mammals (Khakh et al., 2015). In the mammalian cerebellum, there are three types of astroglial cells: Bergmann glial cells (BGs) in the Purkinje cell layer (PCL), astrocytes in the inner granule cell layer (IGL astrocytes), and astrocytes in the white matter (WM astrocytes) (Leto et al., 2016). The cerebellum undergoes significant development after birth. There are two types of astroglial progenitors in the postnatal cerebellum: Bergmann glia-like progenitors (BGLPs) located in the PCL and astrocyte-like progenitors (AsLPs) in the white matter (Joyner & Bayin, 2022). AsLPs are proliferative cells that exhibit a multipolar morphology akin to that of normal astrocytes, and analogous astroglial progenitors are also found in other brain regions. In contrast, BGLPs are proliferative progenitors with unipolar processes that are morphologically indistinguishable from BGs. Such uniquely characteristic astroglial progenitors are rarely observed in other brain regions in mice.

Using a recombination-based lineage tracing method with multiple colors, Cerrato et al. previously suggested that BGLPs at postnatal day 6 (P6) differentiated not only into BGs but also into IGL astrocytes; however, they did not differentiate into WM astrocytes (Cerrato et al., 2018). Due to their morphological similarity, it had been thought that BGLPs could only differentiate into BGs, but their results suggested that BGLPs have a wider differentiation potential than previously imagined. Thus, BGLPs are mysterious cells, but it remains unclear how long they actually exist after birth, what kind of cells BGLPs other than at P6 differentiate into, and what kind of gene expression characteristics BGLPs have.

In this study, we investigated the dynamics of BGLPs, including how their numbers change after birth and how long they persist in the postnatal mouse cerebellum. Next, using the electroporation-based method, we investigated the types of descendant cells that BGLPs at P0 and P6 differentiate into. Furthermore, we successfully extracted the gene expression characteristics of BGLPs at P0 and P6 using image-based spatial transcriptomics (Xenium) on the mouse cerebellum. This study contributes to understanding the astroglial cell development in the postnatal mammalian cerebellum.

## Results

### BGLPs and AsLPs in the postnatal mouse cerebellar development

BGLPs and AsLPs are astroglial progenitors found during the early postnatal stages, but their cell number dynamics and the precise disappearance timing have not been well described. Therefore, we investigated the numbers of the two types of astroglial progenitors, along with BGs and astrocytes, by immunostaining the sagittally-sectioned midline vermis at several postnatal stages in mice (Fig. 1a-c). To identify astroglial lineage cells, we labeled nuclei with SOX9. Cellular morphology was visualized using antibodies against VIMENTIN (at P0-P4) or GFAP (at P6-P21), respectively. KI67 signals were used to discriminate between proliferative progenitors (BGLPs and AsLPs) and postmitotic cells (BGs and astrocytes). BGLPs are SOX9+ and KI67+ cells that localize in the PCL and have unipolar projections extending toward the pia. AsLPs are SOX9+ and KI67+ cells characterized by a multipolar morphology, found in the deeper regions of the cerebellum. We found that the number of BGLPs and AsLPs per section gradually decreased from P0 to P10, and both types of progenitors nearly vanished at P10 (Fig. 1a, b). In contrast, BGs and astrocytes (including IGL and WM astrocytes) increased rapidly from P0 to P10, while little further increase was observed after P10 (Fig. 1a, c). These observations suggest that astroglial progenitors (BGLPs and AsLPs) divide and differentiate into postmitotic astroglial cells (BGs and astrocytes) until P10; however, after this point, these progenitors are rarely present and therefore produce few BGs and astrocytes during normal postnatal cerebellar development.

**Figure 1.**
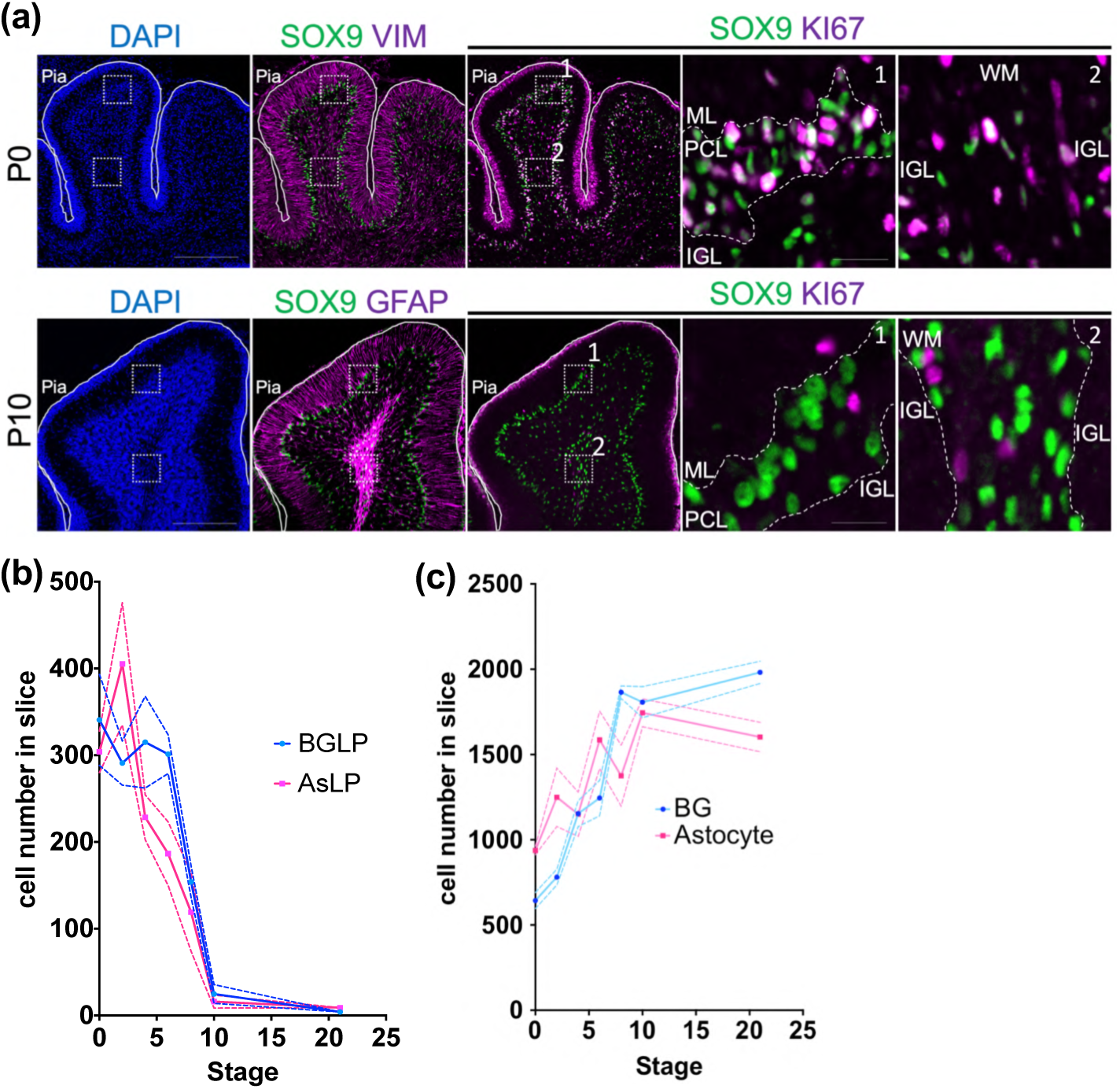
BGLPs and AsLPs in the postnatal mouse cerebellar development. (a) Immunostaining of the P0 and P10 mice cerebellum with DAPI, SOX9, KI67, VIMENTIN and GFAP; Scale bars: 200 μm (left panels), 20 μm (right panel). (b) Number of each astroglial progenitors in P0, P2, P4, P6, P8, P10 and P21 mouse cerebellar sections. Bergmann glia-like progenitor (BGLP) is SOX9 and KI67 double positive and has unipolar fibers (stained with VIMENTIN or GFAP). Astrocyte-like progenitor (AsLP) is SOX9 and KI67 double positive and has multipolar fibers (stained with VIMENTIN or GFAP). (c) Number of each astroglial cells in P0, P2, P4, P6, P8, P10 and P21 mouse cerebellar sections. Bergmann glial cell (BG) is SOX9 positive and KI67 negative, and has unipolar fibers (stained with VIMENTIN or GFAP). Astrocyte is SOX9 positive and KI67 negative and has multipolar fibers (stained with VIMENTIN or GFAP).

### Electroporation method can introduce vectors into BGLPs in the developing cerebellum

Previously, we developed the electroporation (EP) method for introducing DNA vectors into the postnatal mouse cerebellar surface from the pial side (Miyashita et al., 2021; Adachi et al., 2021; Yamashita et al., 2020; Owa et al., 2018) and reported that DNA vectors were introduced not only into GCPs but also into a small number of BG-shaped SOX9+ cells (Owa et al, 2018). To further investigate the cell types of vector introduced cells, we electroporated a CAG-driven mCherry vector (CAG-mCherry) into the cerebellar surface at P0 or P6 and performed immunostaining with cell type-specific markers one day after EP (Fig. 2a, j). When electroporated at P0, about 20 cells per cerebellar lobule were labeled with mCherry at P1 (18.8 ± 1.6, Fig. 2b, f). As expected, most mCherry+ cells belonged to the GC lineage (PAX6+ cells; GCPs and GCs) found in the external granule cell layer (EGL) (92.8 ± 0.3 %, Fig. 2b, g). We also found that this method did not introduce the vector into Purkinje cells (CalbindinD28k+ cells), unipolar brush cells (TBR2+ cells), oligodendrocyte-lineage cells (OLIG2+ cells), or PAX2+ interneurons (Fig. 2b, c, d, g). As previously reported (Owa et al., 2018), approximately 10% (11.8 ± 2.1%) of mCherry+ cells were BG-like cells in the PCL (Fig. 2e, g). These BG-like cells were thought to be BGLPs or BGs, as they expressed SOX9 and VIMENTIN. Because 54.5 ± 3.9% of the mCherry+ BG-like cells were KI67+ (Fig. 2h, i), approximately half of these were BGLPs, while the other half were BGs. The multipolar SOX9+ cells located in the deep cerebellum were rarely labeled with mCherry (Fig. 2g), suggesting that this method does not introduce vectors into AsLPs or astrocytes.

**Figure 2.**
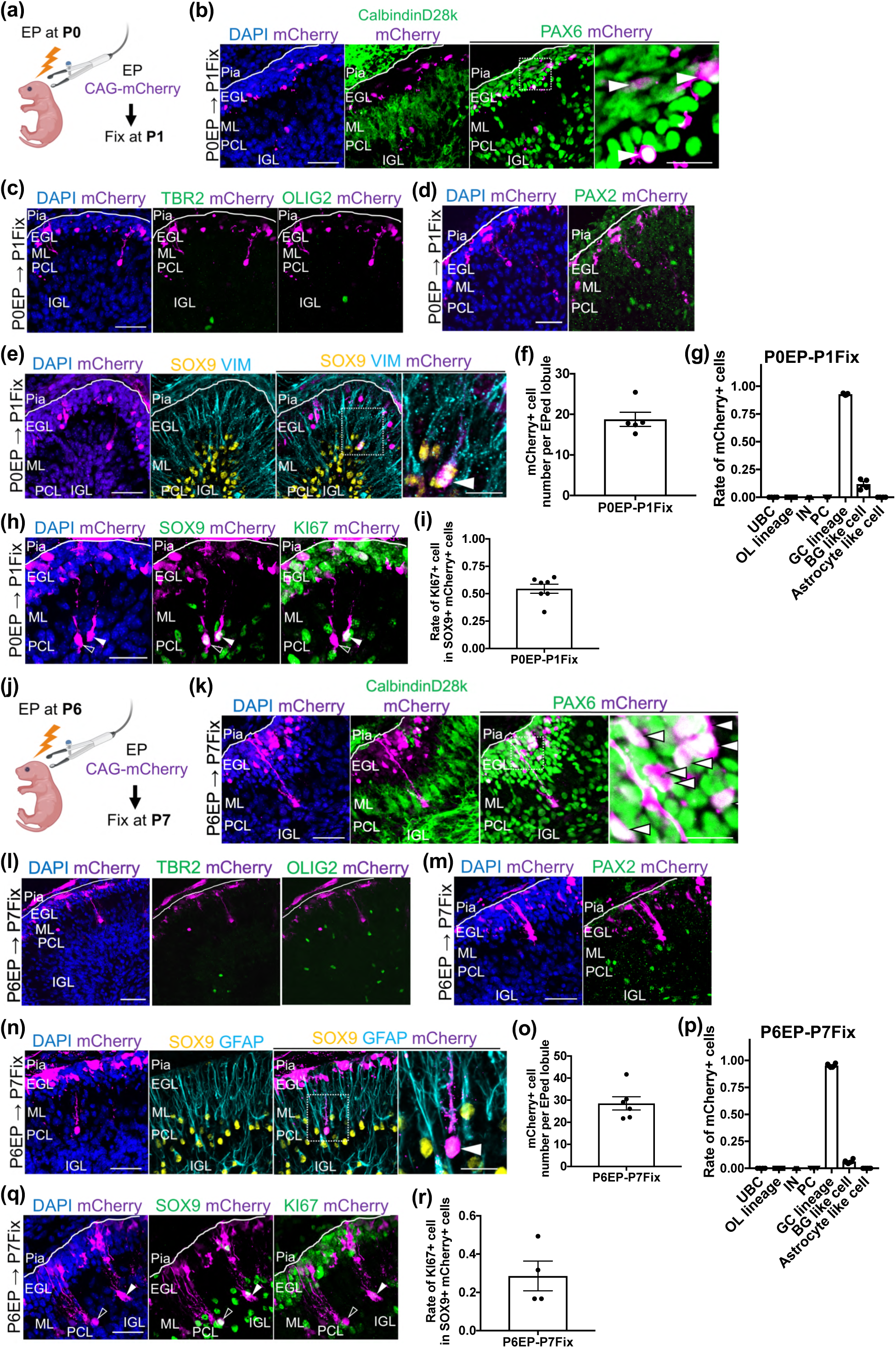
Electroporation method introducing vectors into postnatal BGLPs. (a) Experimental scheme. CAG-mCherry vector was introduced into P0 mouse cerebellar surface by electroporation. Electroporated mouse were fixed at P1. (b-e) Immunostaining with DAPI, RFP (for enhancing mCherry), CalbindinD28K, PAX6, TBR2, OLIG2, PAX2, SOX9 and VIMENTIN in the P1 mouse cerebellum; Scale bars: 50μm for (c), (d) and lower magnification panel of (b) and (e). 30 μm, 10 μm for higher magnification panel of (b) and e), respectively. The white arrowheads in (b) indicate GC lineage cells that are positive for both PAX6 and mCherry which are localized to EGL. The white arrowhead in (e) indicates BG like cell (BGLP or BG) that is positive for both SOX9 and mCherry and has unipolar projections. (f) mCherry+ cell number per electroporated lobule. Animal numbers: *N=5* for the analysis. (g) Unipolar Brush Cell (UBC) is TBR2 positive cell; *N=4*. Oligodendrocyte lineage (OL lineage) is OLIG2 positive cell; *N=4*. Interneuron (IN) is PAX2 positive cell; *N=3*. Purkinje Cell (PC) is CalbindinD28K positive cell; *N=5*. Granule Cell lineage (GC lineage) is PAX6 positive EGL localizing cell; *N=5.* BG like cell is SOX9 positive and has unipolar fibers (stained with VIMENTIN); *N=4*. Astrocyte like cell is SOX9 positive and has multipolar fibers (stained with VIMENTIN); *N=4*. (h) Immunostaining with DAPI, RFP (mCherry), SOX9, KI67 in P1; Scale bars: 40Cμm. White arrowhead indicates BGLP that is positive for SOX9, KI67, and mCherry and has unipolar projections. White open arrowhead indicates BG that is positive for SOX9 and mCherry and has unipolar projections and is negative for KI67. (i) KI67+ cell rate in SOX9+ mCherry+ cells; *N=7*. (j) Experimental scheme. CAG-mCherry vector was introduced into P6 mouse cerebellar surface by electroporation. Electroporated mouse were fixed at P7. (k-n) Immunostaining with DAPI, RFP (mCherry), CalbindinD28K, PAX6, TBR2, OLIG2, PAX2, SOX9 and GFAP in P7 mouse cerebellum; Scale bars: 50μm for (l), (m) and lower magnification panel of (k) and (n). 30 μm and 10 μm for higher magnification panel of (k) and (n), respectively. The white arrowheads in (k) indicate GC lineage cells that are positive for both PAX6 and mCherry which are localized to EGL. The white arrowhead in (n) indicates BG like cell (BGLP or BG) that is positive for both SOX9 and mCherry and has unipolar projections. (o) mCherry+ cell number per electroporated lobule; *N= 6* for the analysis. (p) The markers for UBC, OL lineage, IN, PC and GC lineage used the same antibodies as P1. Animal numbers: *N=4* for UBC, OL lineage, *N=5* for IN, and *N=6* for PC and GC lineage. BG like cell is SOX9 positive and has unipolar fibers (stained with GFAP); *N=4* for the analysis. Astrocyte like cell is SOX9 positive and has multipolar fibers (stained with GFAP); *N=4* for the analysis. (q) Immunostaining with DAPI, RFP (mCherry), SOX9, KI67 in P7; Scale bars: 40Cμm. White arrowhead indicates BGLP that is positive for SOX9, KI67 and mCherry, and has unipolar projections. White open arrowhead indicates BG that is positive for SOX9 and mCherry and has unipolar projections and is negative for KI67. (r) KI67+ cell rate in SOX9+ mCherry+ cells; *N=4* for the analysis. All data are shown as mean ± SEM.

When the mCherry vector was electroporated at P6, we observed about 30 mCherry+ cells at P7 (28.6 ± 2.7, Fig. 2k, o). Of these, approximately 5% were BG-like cells (6.3 ± 0.6 %, Fig. 2n, p), while 95% were GC lineage cells (GCPs and GCs) localized to the EGL (95.3 ± 0.5 %, Fig. 2k, p). We did not detect mCherry signals in other cell types within the cerebellum (Fig. 2k, l, m, p). Immunostaining with KI67 further indicated that roughly 30% of the mCherry+ BG-like cells were BGLPs (28.6 ± 6.7 %, Fig. 2q, r). We did not observe SOX9+ multipolar cells (AsLPs or astrocytes) in the deep cerebellum (Fig. 2p). These findings suggest that this EP method can introduce vectors into BGLPs and BGs, but not into AsLPs or astrocytes.

### Differentiation of P0 BGLPs and P6 BGLPs into cerebellar astroglial subtypes

Next, we sought to investigate which types of astroglial cells (BGs, IGL astrocytes, and WM astrocytes) are generated from BGLPs during postnatal cerebellar development. We demonstrated that our EP can introduce the CAG-mCherry vector into BGLPs and BGs, as well as GC-lineage cells (GCPs and GCs), but not into any other cell types at postnatal stages (Fig. 2). Previous cell lineage analyses revealed that GC-lineage cells give rise only to GCs and do not differentiate into other cell types during normal cerebellar development (Vladoiu et al., 2019; Machold & Fishell, 2005; Wang et al, 2005). This suggests that in the CAG-mCherry EP method, the mCherry/SOX9-double positive cells after EP are thought to be the electroporated BGs themselves or descendant astroglial cells derived from the electroporated BGLPs.

We electroporated the CAG-mCherry into the cerebellum at P6 and fixed the samples for immunostaining at P10 or P30 (Fig. 3a, b). Since BGLPs and AsLPs are nearly absent at P10 (Fig. 1), most SOX9+ cells should be BGs, IGL astrocytes, or WM astrocytes at P10 and P30. We found that the numbers of mCherry/SOX9-double positive cells per lobule were 1.8 ± 0.3 at P10 and 3.4 ± 0.6 at P30 (Fig. 3c). In the immunostained cerebella at P10 and P30, mCherry/SOX9-double positive cells were classified into BGs, IGL astrocytes, and WM astrocytes based on their morphology and positions. We observed that among mCherry/SOX9-double positive cells, 64.3 ± 6.0% of cells were BGs and 34.6 ± 6.2% were IGL astrocytes in P10 cerebella. Similarly, 78.1 ± 5.7% were BGs and 21.4 ± 5.2% were IGL astrocytes in P30 cerebella (Fig. 3d). We found no WM astrocytes among mCherry/SOX9-double positive cells. This suggests that BGLPs at P6 differentiate into BGs and IGL astrocytes, but not into WM astrocytes, during cerebellar development. This result aligns with the previous finding obtained by a recombination-based lineage tracing method (Cerrato et al., 2018), indicating that our electroporation-based method is a reliable tool for investigating BGLP progeny cells.

**Figure 3.**
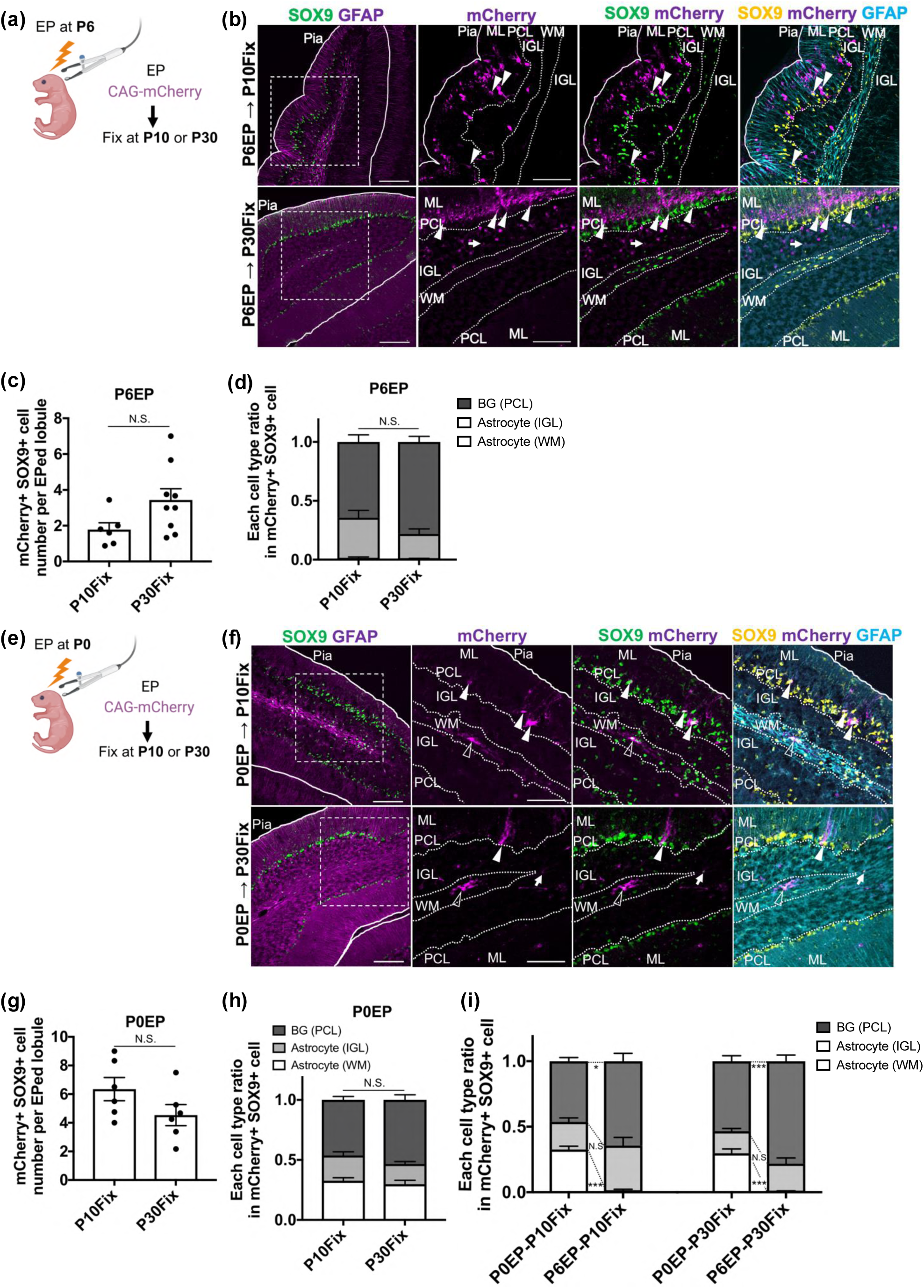
P0 mouse BGLPs differentiate into WM astrocytes, whereas P6 mouse BGLPs do not. (a) Experimental scheme. CAG-mCherry vector was introduced into P6 mouse cerebellar surface by electroporation. Electroporated mouse were fixed at P10 or P30. (b) Immunostaining with SOX9, GFAP, RFP (mCherry) in P10 and P30; Scale bars: 100 μm (left panels) and 70Cμm (right panels). White arrowhead indicates BG that is positive for SOX9, and mCherry, and has unipolar projections. White arrow indicates IGL astrocyte that is positive for SOX9, and mCherry, has multipolar projections, and localized in IGL. (c) mCherry+ SOX9+ cell number per electroporated lobule; *N=6* for P6EP-P10Fix and *N=9* for P6EP-P30Fix. Statistical tests were performed by Student’s *t* test; *p* values were represented as; N.S. for *p* > 0.05. (d) Distribution of mCherry+ astroglial cells; *N=6* for P6EP-P10Fix and *N=9* for P6EP-P30Fix analysis. Statistical tests were performed by Sidak’s multiple comparisons test; *p* values were represented as; N.S. for *p* > 0.05. (e) Experimental scheme. CAG-mCherry vector was introduced into P0 mouse cerebellar surface by electroporation. Electroporated mouse were fixed at P10 or P30. (f) Immunostaining with SOX9, GFAP, RFP (mCherry) in P10 and P30; Scale bars: 100 μm (left panels) and 70 μm (right panels). White arrowhead indicates BG that is positive for SOX9, and mCherry, and has unipolar projections. White arrow indicates IGL astrocyte that is positive for SOX9, and mCherry, has multipolar projections, and localized in IGL. White open arrowhead indicates WM astrocyte that is positive for SOX9, and mCherry, has multipolar projections, and localized in WM. (g) mCherry+ SOX9+ cell number per electroporated lobule; *N=6* for both stages. Statistical tests were performed by Student’s *t* test; *p* values were represented as; N.S. for *p* > 0.05. (h, i) Distribution of mCherry+ astroglial cells; *N=6* for P0EP-P10 Fix, P6EP-P10Fix and P0EP-P30Fix, and *N=9* for P6EP-P30Fix; N.S > 0.05, \**p* <0.05, \*\*\**p* < 0.001, Sidak’s multiple comparisons test. All data are shown as mean ± SEM.

Next, we electroporated CAG-mCherry into the cerebellum at P0 and fixed the samples for immunostaining at P10 or P30 (Fig. 3e, f). We found that the numbers of mCherry/SOX9-double positive cells per lobule were 6.3 ± 0.7 at P10 and 4.4 ± 0.7 at P30 (Fig. 3g). Surprisingly, we observed some mCherry/SOX9-double positive cells localized in the WM (i.e., WM astrocytes). Among those double positive cells, 46.2 ± 2.9%, 21.1 ± 2.9%, and 32.7 ± 2.5% were BGs, IGL astrocytes, and WM astrocytes in the electroporated cerebella at P10, while 53.3 ± 4.3%, 17.0 ± 1.9%, and 29.7 ± 3.4% were BGs, IGL astrocytes, and WM astrocytes at P30 (Fig. 3f, h, i). These observations suggest that BGLPs at P0 differentiate not only into BGs and IGL astrocytes but also into WM astrocytes, further implying that BGLPs at P0 may have wider differentiation potential than those at P6.

### Electroporation experiment using hGFAP-EGFP vector

The observation that BGLPs at P0 produced not only BGs and IGL astrocytes but also WM astrocytes was somewhat surprising, as it differed from the previous report on P6 BGLPs (Cerrato, et al., 2018). Therefore, we attempted to confirm this result using a different method. The hGFAP promoter is known to specifically promote gene transcription in astroglial progenitors and astroglial cells in the postnatal cerebellum (Zhuo et al., 1997). We generated the hGFAP-EGFP vector, in which EGFP is designed to be expressed under the control of the hGFAP promoter. We co-electroporated hGFAP-EGFP and CAG-mCherry vectors into the cerebellum of P0 mice. One day after EP, we observed 1–2 EGFP+ cells per lobule (1.4 ± 0.1, Fig. 4a-c). SOX9 and VIMENTIN immunosignals, cell positions, and cell morphologies indicated that most EGFP+ cells (94.8 ± 3.3%) were BGLPs or BGs (Fig. 4b, d). Immunostaining with KI67 showed that 93.3 ± 0.4% of those BG-like cells were BGLPs (Fig. 4e, f). In contrast, EGFP+ cells did not express NEUROD1, indicating that this hGFAP-EGFP hardly labeled GCs (Fig. 4g). This suggests that hGFAP-EGFP electroporation at P0 can specifically label BGLPs but not GC-lineage cells.

**Figure 4.**
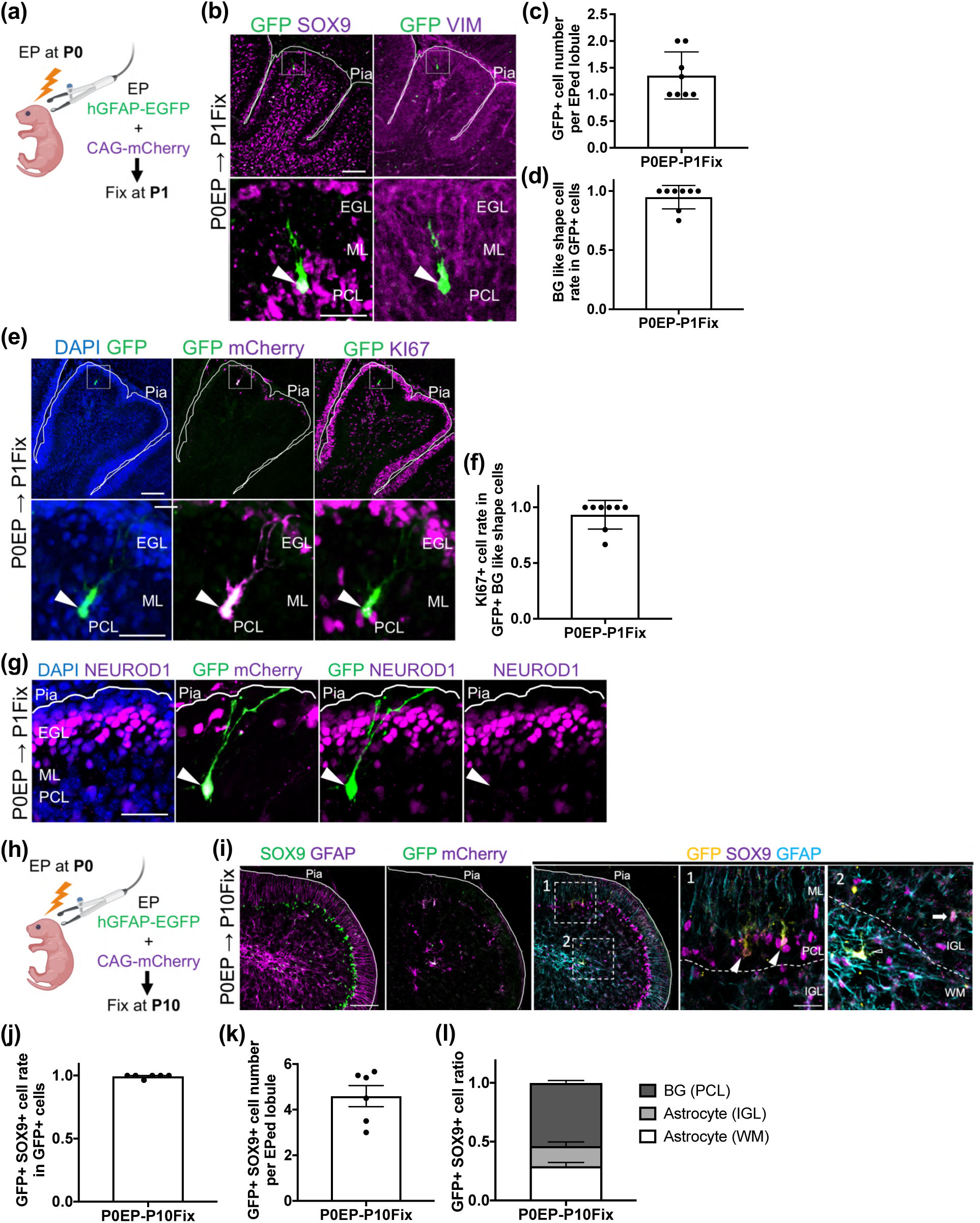
P0 mouse BGLPs specifically labeled with hGFAP-EGFP give rise to WM astrocytes. (a) Experimental scheme. hGFAP-EGFP vector and CAG-mCherry vectors were introduced into mouse cerebellum at P0 and fixed at P1. (b) Immunostaining with SOX9, VIMENTIN, GFP at P1; Scale bars: 100 μm (upper panels) and 50 μm (lower panels). (c) GFP+ cell number per electroporated lobule; *N=8* for the analysis. (d) BG-like shape cell rate in GFP+ cells; *N=8* for the analysis. (e) Immunostaining with DAPI, GFP, RFP (mCherry), KI67 at P1; Scale bars: 100 μm (upper panels) and 50 μm (lower panels). (f) KI67+ cell rate in GFP+ BG-like shape cells; *N=8* for the analysis. (g) Immunostaining with DAPI, GFP, RFP (mCherry), NEUROD1 at P1; Scale bars: 50 μm. (h) Experimental scheme. hGFAP-EGFP and CAG-mCherry vectors were introduced into mouse cerebellum at P0 and fixed at P10. (i) Immunostaining with SOX9, GFP, RFP (mCherry), GFAP at P10; Scale bars: 100 μm (left panels) and 40 μm (right panels). White arrowhead indicates BG that is positive for SOX9, and GFP, and has unipolar projections. White arrow indicates IGL astrocyte that is positive for SOX9, and GFP, has multipolar projections, and localized in IGL. White open arrowhead indicates WM astrocyte that is positive for SOX9, and GFP, has multipolar projections, and localized in WM. (j) GFP+ SOX9+ cell rate in GFP+ cells; *N=6* for the analysis. (k) GFP+ SOX9+ cell number per electroporated lobule; *N=6* for the analysis. (l) Distribution of GFP+ astroglial cells; *N=6* for the analysis. All Data are shown as mean ± SEM.

To investigate the progeny cells of P0 BGLPs, the cerebella of P0 mice were electroporated with hGFAP-EGFP and CAG-mCherry, followed by immunostaining with several antibodies at P10 (Fig. 4h, i). In these P10 samples, the proportion of astroglial cells (SOX9+ cells) among EGFP+ cells was nearly 100% (99.4 ± 0.5, Fig. 4i, j). We observed approximately 4.6 ± 0.4 astroglial cells (SOX9+ cells) labeled with EGFP per lobule (Fig. 4i, k). EGFP+ astroglial cells (SOX9+ cells) were found in the PCL, IGL, and WM. Based on their morphologies (visualized with GFAP) and positions, these cells were thought to be BGs, IGL astrocytes, and WM astrocytes, respectively (53.7 ± 2.0%, 17.1 ± 3.5%, and 29.2 ± 2.9%) (Fig. 4i, l), which was generally consistent with the results of the CAG-mCherry electroporation experiment (Fig. 3f, h, i). From these results, it was confirmed that P0 BGLPs produce all types of astroglial cells (BGs, IGL astrocytes, and WM astrocytes) in the cerebellum.

Next, we electroporated hGFAP-EGFP and CAG-mCherry into the P6 mouse cerebellum and fixed the samples at P7, one day after EP (Supplemental Fig. 1a). Unexpectedly, some GCP/GC-like cells in the EGL were labeled with EGFP (74.5 ± 7.2% of GFP+ cells), in addition to BG-like cells (BGLPs and BGs) (Supplemental Fig. 1b-d). We thought that the hGFAP promoter in the vector induced gene expression very specifically in astroglial cells at P0/P1, but this specificity might become somewhat ambiguous at later stages. Therefore, the experiment using the hGFAP-EGFP vector was conducted solely to analyze P0 BGLPs.

### Spatial transcriptomic analysis reveals distinct molecular signatures of P0 and P6 BGLPs

To investigate the differences in the molecular properties of BGLPs at distinct stages, we conducted image-based spatial transcriptomics on Xenium platform using cerebellar sections from P0 and P6 mice (Fig. 5). In total, 5,146 and 19,892 cell data were obtained in the P0 and P6 mouse cerebellar slices, respectively. We applied unsupervised clustering, which identified 24 unique clusters of cells shared by the two stages (Fig. 5a, b, Supplemental Fig. 2a). We annotated each cluster with the expression of cell markers, and cell morphologies and locations within the cerebellum (Fig. 5c, Supplemental Fig. 3). We found that clusters 6 (1,117 cells), 11 (820 cells), and 13 (607 cells) corresponded to P0 and P6 astroglial lineage cells. To further characterize the astroglial lineage cells, we extracted astroglial lineage cells of clusters 6, 11, and 13, and performed sub-clustering, ultimately obtaining 10 sub-clusters. (Fig. 5d, Supplemental Fig. 2b). Most cells in sub-clusters 0, 6, and 7 were derived from P0, while sub-clusters 1, 2, 3, 4, and 5 originated from P6 (Supplemental Fig. 2b). Sub-clusters 8 and 9 were excluded from further analysis due to their small cell populations and dispersed localization (Supplemental Fig. 2c). Based on the expression profiles of known BG-like cell-specific markers (*Gdf10*, *Gria1*, *Sept4*) and proliferative markers (*Pcna*, *Mki67*), along with spatial information, it was indicated that sub-cluster 7 was BGLPs in P0 mouse (Fig. 5e, f, Supplemental Fig. 2d-h), while sub-clusters 1 and 4 were BGLPs in P6 mouse (Fig. 5e, g, Supplemental Fig. 2i-m).

**Figure 5.**
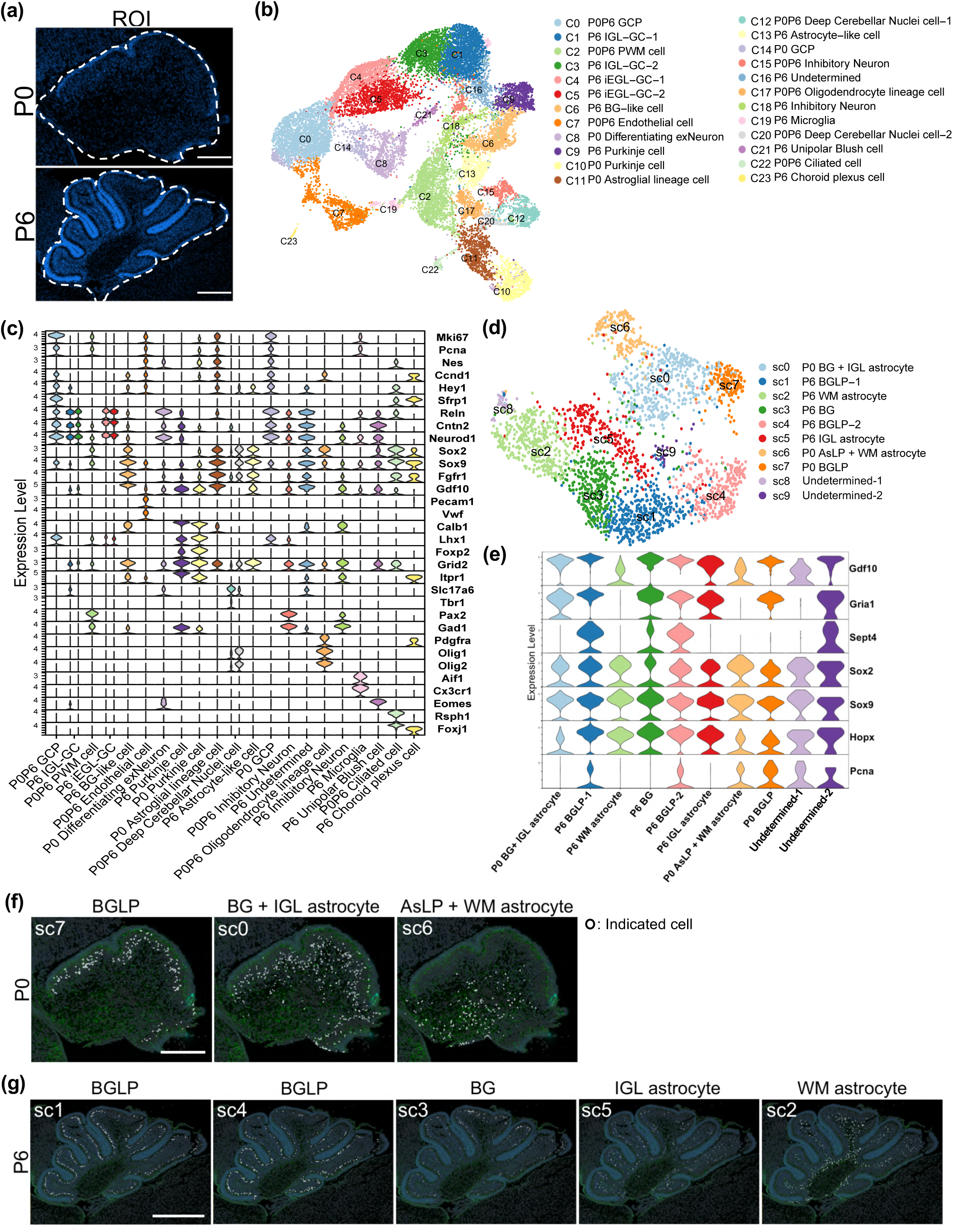
Spatial transcriptomic analysis identifies BGLP clusters in P0 and P6 mouse cerebellum. (a) Cerebellar slices used for spatial transcriptomics analysis. The panels show region of interests (ROI) which was stained with DAPI. Scale bars: 200 μm for P0 and 500 μm for P6. (b) Uniform manifold approximation and projection (UMAP) dimensional reduction was performed using segmented cells and 24 distinct clusters were identified. (c) Gene expression profiles of each cerebellar cell markers across the 24 identified clusters. Cluster annotation was performed based on the expression profiles of cell markers and spatial localization information. (d) UMAP dimensional reduction was performed using P0 and P6 astroglial lineage cell clusters (Cluster 6, 11, 13). 10 distinct sub-clusters were identified. (e)-(g) Annotation of each astroglial lineage cell sub-cluster. Annotation was performed based on the expression profiles of BG-like cell specific markers and proliferative marker (e), and spatial localization information (f, g). Scale bars: 200 μm for (f) and 500 μm for (g).

Next, we compared the expression profiles between the BGLP sub-clusters of P0 (sub-cluster 7) and P6 (sub-clusters 1 and 4) (Fig. 6a). We identified 311 differentially expressed genes (DEGs). Of these, 224 genes were more highly expressed in the BGLP sub-cluster at P0 compared to P6 (referred to as ‘upregulated genes in P0 BGLP’), while 87 genes showed higher expression at P6 than at P0 (referred to as ‘upregulated genes in P6 BGLP’). (Average Log2 Fold Change > 1, p_val_adj < 0.05, Fig. 6a, Table 1). We then carried out the gene ontology (GO) analysis on identified up- and down-regulated DEGs (Fig. 6b, c). Several GO categories for “upregulated genes in P0 BGLP” were associated with cell cycle progression, including “nuclear division,” “mitotic cell cycle phase transition,” and “mitotic nuclear division” (Fig. 6b). Interestingly, the “upregulated genes in P0 BGLP” included *Nes*, *Sox4*, and *Sox11*, known as stem cell markers (Fig. 6d). On the other hand, some GOs for “upregulated genes in P6 BGLP” implicated glial cell differentiation, such as “gliogenesis,” “glial cell differentiation,” and “astrocyte differentiation” (Fig. 6c). The “upregulated genes in P6 BGLP” included BG-like cell-specific markers such as *Gdf10*, *Gria1*, *Tnc*, and *Sept4* (Fig. 6e). These findings suggest that BGLPs in P0 mice exhibit more stem cell-like characteristics, whereas BGLPs in P6 mice display more BG-like features. These results are consistent with our *in vivo* observations, which indicate that BGLPs in P0 mice demonstrate higher differentiation ability than those in P6 mice (Fig 6g).

**Figure 6.**
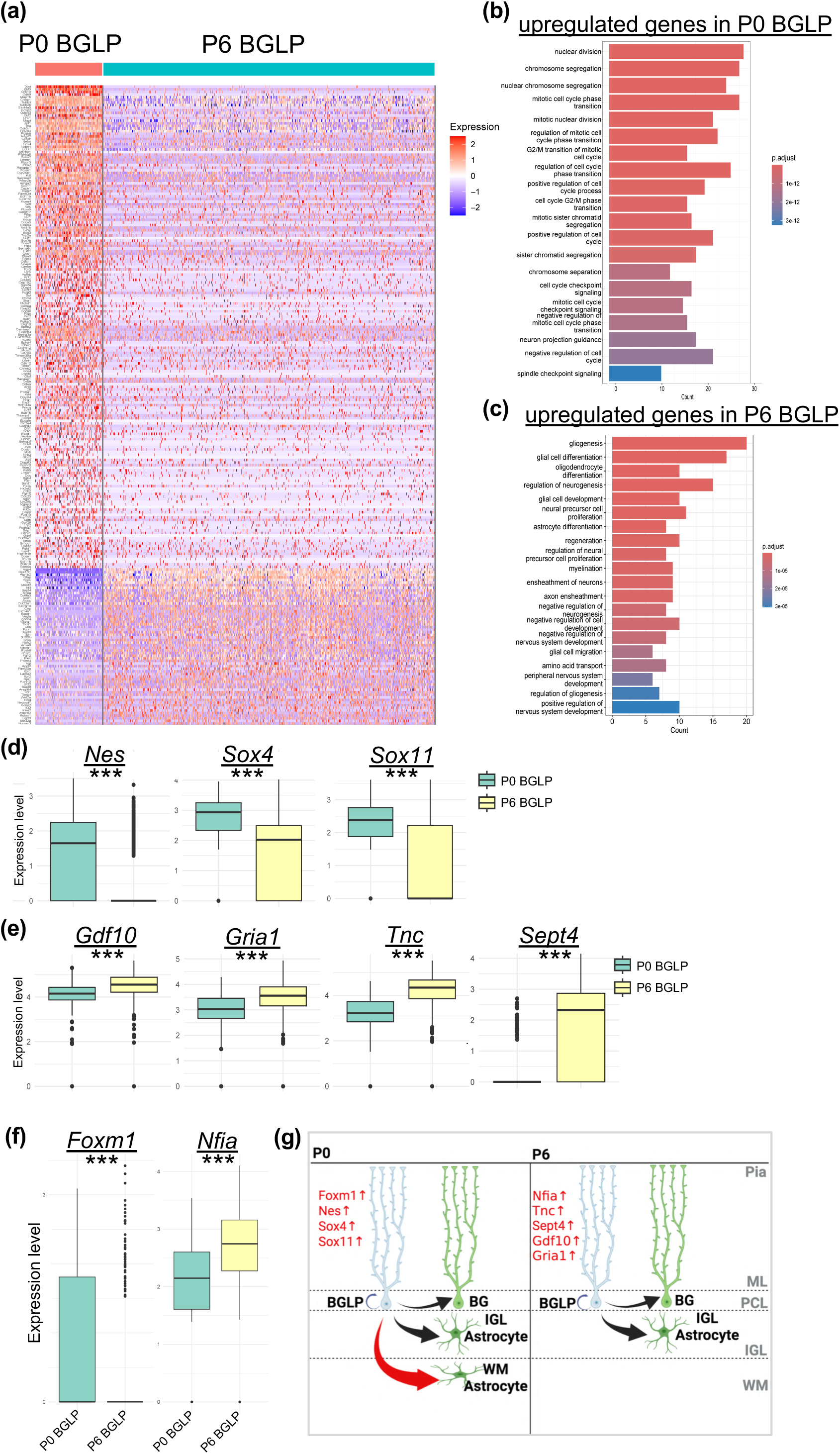
P0 mouse BGLPs show higher stem cell marker expression, while P6 mouse BGLPs exhibit elevated BG-like cell specific gene levels. (a) Expression profiles of 311 DEGs between BGLP from P0 and P6 mouse were visualized by heatmap. 224 genes were upregulated in BGLPs at P0, while 87 genes were upregulated in BGLPs at P6. Due to the difference in sex between the P0 and P6 mouse, the identified differentially expressed genes (DEGs) included sex-linked genes, such as *Xist*. (b) GO analysis was performed for the 224 upregulated genes in BGLPs at P0. (c) GO analysis was performed for the 87 upregulated genes in BGLPs at P6. (d) Box plots of *Nes*, *Sox4* and *Sox11*, genes associated with the maintenance of pluripotency, \*\*\**p* < 0.001. (e) Box plots of BG-like cell specific markers, *Gdf10*, *Gria1*, *Tnc* and *Sept4*. \*\*\**p* < 0.001. (f) Box plots of *Foxm1* and *Nfia*, putative upstream regulators of DEGs in BGLPs at P0 and BGLPs at P6, \*\*\**p* < 0.001. (g) A schematic model illustrating the differences in differentiation potential and gene expression profiles of BGLPs between P0 and P6 mice. At P0, BGLPs give rise not only to BGs and IGL astrocytes, but also to WM astrocytes. Compared to BGLPs in P6 mouse, BGLPs in P0 mouse exhibit higher expression of genes associated with stemness. In contrast, BGLPs in P6 mouse show elevated expression of BG-like cell–specific genes.

**Table 1.** Differentially Expressed Genes in P0 and P6 BGLPs Gene lists represent differentially expressed genes in BGLPs at postnatal day 0 (P0, Sheet1) and P6 (Sheet2) as identified by image-based spatial transcriptomic analysis on Xenium platform.

**Table 2.** Top five “upstream candidates for P0 BGLP” and “upstream candidates for P6 BGLP” predicted from Enrichr analysis Top five “upstream candidates for P0” and “upstream candidates for P6” by Enrichr using the databases (“ChEA2022”, “ENCODE and ChEA Consensus TFs from ChIP-X”, “ARCHS4 TFs Coexp”, “TF Perturbations Followed by Expression”, “TRRUST Transcription Factors 2019”, “Enrichr Submissions TF-Gene Coocurrence”, “Transcription Factor PPIs”).

Finally, we explored the upstream transcription factors (TFs) that potentially regulate the gene expression of BGLPs at P0 and P6. We utilized Enrichr (Chen et al., 2013) to examine TFs that could serve as upstream regulators of the “upregulated genes in P0 BGLP” and “upregulated genes in P6 BGLP” (referred to as “upstream candidates for P0 BGLP” and “upstream candidates for P6 BGLP,” respectively) (Table. 2). Interestingly, *Foxm1* was included in both “upstream candidates for P0 BGLP” and “upregulated genes in P0 BGLP” (Fig. 6f). Similarly, *Nfia* was found in both “upstream candidates for P6 BGLP” and “upregulated genes in P6 BGLP” (Fig. 6f). This implies that *Foxm1* and *Nfia* might be involved in regulating the gene expression profiles of BGLPs at P0 and P6, respectively (Fig. 6g).

## Discussion

BGLPs are unique astroglial progenitors with a unipolar shape that divide after birth to generate astroglial cells. In the past, a few studies related to BGLPs have been reported. Li et al. demonstrated that certain BG-like cells in the PCL of the postnatal mouse cerebellum are proliferative (Li et al, 2013), and these cells are later called BGLPs. Wojcinski et al. reported that, when the cerebellum suffered significant damage from radiation exposure, BGLPs contributed to the recovery of cerebellar structure and function by generating not only astroglial cells but also GCPs (Wojcinski et al, 2017). However, in normal development, Cerrato et al. showed that, by recombination-based lineage tracing, BGLPs produce only astroglial cells, but not GCPs (Cerrato et al., 2018).

This indicates that BGLPs acquire wider differentiation potential only in emergency situations, such as irradiation, but only produce astroglial lineage cells during normal development. Furthermore, they demonstrated that P6 BGLPs produce BG and IGL astrocytes but not WM astrocytes. Despite these previous works, many aspects of BGLPs still remain unclear, including their cell number dynamics, the types of cells they produce, and their gene expression profiles during postnatal development.

In this study, we observed that BGLPs decreased from P0 to P10 and nearly vanished by P10. Our in vivo electroporation experiment in the normally developing cerebellum showed that BGLPs at P0 produce BGs, IGL astrocytes, and WM astrocytes, while P6 BGLPs generate BGs and IGL astrocytes but not WM astrocytes. This indicates that P0 BGLPs have a wider potential for cell differentiation than previously believed (Cerrato et al., 2018). To further investigate the molecular nature of BGLPs, we used image-based spatial transcriptomics (Xenium 5k) on mouse cerebella at postnatal stages (P0 and P6). To our knowledge, this is the first report of image-based spatial transcriptomics with single-cell resolution in the cerebellum of mice during the postnatal developmental period. We successfully distinguished different types of astroglial progenitors and astroglial cells into separate clusters based on their gene expression profiles and spatial locations. Previously, single-cell RNA-seq (scRNA-seq) has been used to attempt the clustering of astroglial progenitors and astroglial cells in the postnatal cerebellum (Bayin et al., 2021, Mockenhaupt et al., 2023). However, in conventional scRNA-seq, it seemed difficult to properly cluster each type of progenitors and astroglial cells because the spatial information was lost. We have successfully extracted transcriptomic profiles of BGLPs in the postnatal cerebellum, which have not been studied much to date, by performing image-based spatial transcriptome analysis. Since this transcriptome data includes information on all cell types in the postnatal developing cerebellum, it will undoubtedly contribute to overall cerebellar development research.

## Experimental procedures

### Animals

All animal experiments were approved by the Animal Care and Use Committee of the National Institute of Neuroscience, National Center of Neurology and Psychiatry (Tokyo, Japan; Project 202017R2). ICR mice, purchased from SLC were used in all animal experiments. Mice were raised under SPF conditions, with a 12-h light/dark cycle, and were allowed to freely eat and drink food and water. Both male and female mice were used in this study.

### Plasmids

The hGFAP-EGFP vector was constructed by removing iCre from the purchased hGFAP promoter-driven EGFP-T2A-iCre vector (VB1131; Vector Biolabs) using restriction enzyme digestion. CAG-mCherry was provided by Dr. N. Masuyama.

### In vivo electroporation

*In vivo* electroporation of postnatal mice has been previously described (Adachi et al., 2021). Briefly, expression plasmids were diluted to 1 μg/μL with Milli-Q water. Fast Green was added to achieve a concentration of 0.02% to visualize the plasmid solution. The pups were anesthetized on ice before electroporation, and 10 μL of plasmid solution was injected onto the surface of the cerebellar vermis region of P0 or P6 pups. For P0 and P6, electrical pulses of 70 V and 80 V, respectively, were used for electroporation. The other conditions for the electric pulse were the same for both stages (pulse length of 50 ms, pulse interval of 150 ms, and seven times). Forceps-type electrodes (CUY650P5-3, NEPA) were used for electroporation. After electroporation, the pups were kept warm at 37 °C until they recovered and then returned to their litter. In this experimental method, genes were mainly introduced into lobules IV–VI of the cerebellar vermis. During electroporation of hGFAP-EGFP, CAG-mCherry was co-introduced for normalization purposes.

### Immunohistochemistry and antibodies

Mice less than 10 days old were decapitated, and their brains were collected and shaken while immersed in 4% PFA solution overnight. Mice older than 10 days were perfused and fixed, and their brains were collected and shaken while immersed in a 4% PFA solution overnight. The brains soaked in 4% PFA, were then soaked in 1×PBS, 10% sucrose, and 30% sucrose overnight and shaken. The brains were then embedded in an O.C.T compound (Sakura Finetek). Frozen brains were sagittally sectioned at a thickness of 16 μm using a cryostat (CM3050 S; Leica). Cryosections were washed twice with PBS for 10 min and then incubated at room temperature with 10% normal donkey serum (S30-100ML; Millipore) containing 0.2% PBST (Triton) for 40 min. After blocking, the sections were incubated with primary antibodies diluted with blocking solutions at 4 °C for 16 h. The following primary antibodies were used: mouse anti-CalbinidinD28K (1:500; C9848; Sigma-Aldrich), rat anti-GFAP (1:500; AB5541; Merck Millipore), chicken anti-GFP (1:1000; ab1218; Abcam), rat anti-Ki67 (1:500; 14-5698-82; Invitrogen), goat anti-NeuroD1 (1:500; AF2746; R&D), goat anti-Olig2 (1:500; AF2418; R&D Systems), rabbit anti-PAX2 (1:500; 71-6000; Invitrogen), rabbit anti-PAX6 (1:500, 901301; Biolegend), rabbit anti-RFP (1:500; PM005; MBL), rat anti-RFP (1:1000; 5F8-20; Proteintech Group, Inc.), goat anti-Human SOX9 (1:500; AF3075; R&D Systems), rabbit anti-Tbr2/Eomes (1:1000; ab183991; Abcam), and rabbit anti-Vimentin (1:500; D21H3; Cell Signaling). Subsequently, slides were rinsed with PBS (10 min, twice) and incubated with secondary antibodies conjugated with Alexa Fluor 405, Alexa Fluor 488, Alexa Fluor 568, Alexa Fluor 647 (1:400; Abcam), or Alexa Fluor 488 (1:400; Jackson) and DAPI (25 μg/mL; Invitrogen) in 0.2% PBST at room temperature for 2 h. The slides were rinsed again with PBS (10 min, twice) and mounted with ProLong Glass Antifade Mountant (P36984; LTJ).

### Image acquisition and quantification

The entire sagittal section of the cerebellar vermis was observed under a microscope and analyzed to examine the distribution of astroglial cells in mice at each stage of the experiment. In the experiment introduced genes by electroporation, lobules IV to VI of the cerebellar vermis were observed under a microscope and analyzed. At least two images were obtained from each participant. Images were obtained using a confocal microscope, SpinSR10 (Olympus). The acquired images were adjusted and analyzed using ImageJ software (RSB). All graphs were generated using GraphPad Prism software (GraphPad Software).

### Xenium experiment

FFPE brain samples from ICR P0 and P6 mice were sectioned into 5 μm slices and placed onto Xenium slides. The Xenium Prime 5 K Mouse Pan Tissue and Pathways Panel, which targets 5001 genes, was used for the experiment. Deparaffinization, decrosslinking, probe hybridization, ligation, and signal detection were performed according to the manufacturer’s protocol. Cell segmentation and transcript counting were automatically performed using the Xenium Analyzer.

### Xenium data processing

Regions of interest (ROIs) covering the entire cerebellum of P0 and P6 mice were manually selected using Xenium Explorer (version). The ROIs contained 5,146 cells from P0 mouse and 19,892 cells from P6 mouse. Cell ID information is provided in the supplementary file.

Data analysis was conducted in R (v4.4.2) using the Seurat package (v5.2.1). Gene expression data from the P0 and P6 cerebellum were merged and filtered using the criteria (nCount > 300 and nFeature > 300). After quality filtering, log normalization, dimensionality reduction, and clustering were performed using Seurat functions with default parameters. This analysis resulted in the identification of 24 clusters. Each cluster was annotated based on marker gene expression(reference) and spatial information. Clusters 6, 11, and 13, identified as astroglial lineage clusters, were extracted for subclustering. To remove the influence of cell cycle-related gene expression, cell cycle scoring and regression were performed (Nestorowa et all., 2016), followed by dimensionality reduction and clustering, which yielded 10 sub-clusters. Among these, sub-clusters 7 (BGLPs at P0) and 1 and 4 (BGLPs at P6) were identified as BGLP clusters based on marker gene expression, cell morphology, and spatial localization. Differential expression analysis between BGLP clusters from the P0 and P6 cerebellum was performed using Seurat’s FindMarkers function. Differentially expressed genes (DEGs) were defined by an adjusted p-value < 0.05 and log_₂_ fold change. Gene Ontology (GO) enrichment analysis was performed on the DEGs using the clusterProfiler package (v4.14.4), focusing on the top 20 categories of the “Biological Process” ontology. A heatmap was generated using DEGs with an average log_₂_ fold change > 1 and adjusted p-value < 0.05. To identify potential upstream transcription factors potentially regulating the expressions of DEGs, enrichment analysis was conducted using the enrichR package (Chen et al., 2013) separately for the 224 and 88 DEGs. All codes for generating results are available in the Hoshino Lab GitHub repository.

### Statistical analyses

Individual animals or trials were considered as biological replicates. All mice were prepared under identical experimental conditions. All data are presented as mean ± SEM. All statistical analyses were performed using GraphPad Prism software (GraphPad Software). Statistical tests were performed using Student’s *t* tests and Sidak’s multiple comparison tests; *p* values are represented as N.S. for *p*C>C0.05; \**p*C<C0.05, \*\**p*C<C0.01, and \*\*\**p*C<C0.001.

## Acknowledgements

This work was supported by JSPS KAKENHI (Grant Numbers JP22H02730 to MH, 21K20853 and 23K14203 to TA); AMED (Grant Numbers 24wm0425005h0004 and 24ek0109764h0001 to MH), an Intramural Research Grant of NCNP (4–5, 4-6, 6-9 to MH); Japan Health Research Promotion Bureau (JH) under Research Fund (2020-B-07 and 2024-D-01 to MH); Multilayered Stress Diseases (JPMXP1323015483 to MH); JST SPRING, (Grant Numbers JPMJSP2180 to KS); Tokumori Yasumoto Memorial Trust (MH) and Takeda Science Foundation (Grants 2024049458 to TA). We would like to thank Editage (www.editage.jp) for English language editing.

## Author contributions

T.A. and M.H. supervised this project. K.S., T.A., T.O. and M.H. designed the study. K.S., T.A. and M.H. wrote the manuscript. K.S., T.A., E.I., I.H., M.M., J.K., K.N., S.M. and T.O. performed experiments. K.S., T.A., I.H., S.M. and T.O. analyzed data.

## Data availability

All data used in this study is available upon request.

## Code availability

The custom code associated with this study is available at https://github.com/Hoshino-lab/Suyama_Cerebellum_5K.

---

**Supplemental Figure 1.**
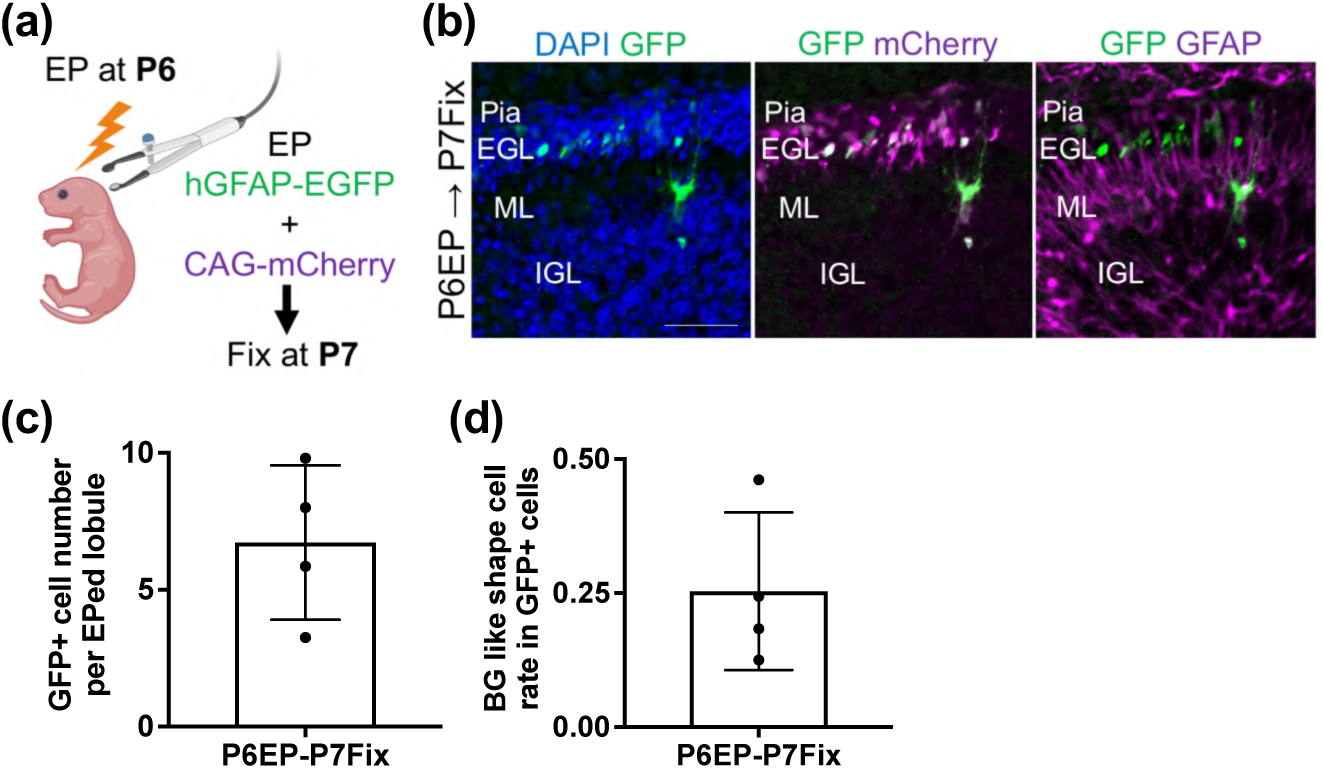
Electroporation of hGFAP-EGFP into the cerebellar surface of P6 mice. (a) Experimental scheme. The hGFAP-EGFP and CAG-mCherry vectors were introduced into the mouse cerebellum at P6 and fixed at P7. (b) Immunostaining with DAPI, GFP, RFP (mCherry), GFAP in P7; Scale bars: 70 μm. (c) GFP+ cell number per electroporated lobule; *N=4* for the analysis. (d) BG-like cells in GFP+ cells; *N=4* for analysis. All data are shown as mean ± SEM.

**Supplemental Figure 2.**
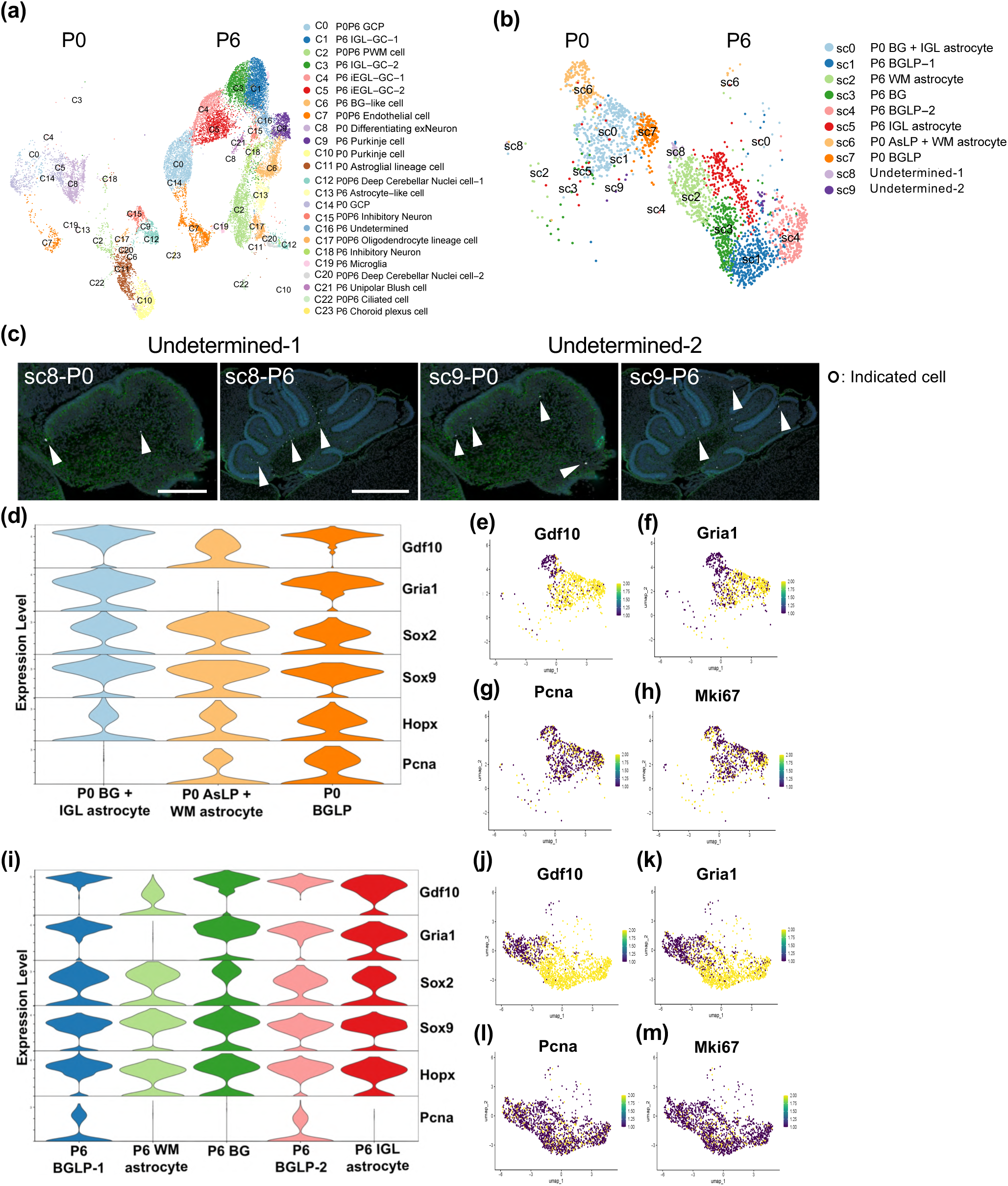
Supplemental data for Figure 5. (a) The 24 identified clusters visualized separately for each stage. (b) The 10 identified sub-clusters visualized separately for each stage. (c) Spatial localization information of sub-cluster 8 and 9. Scale bars: 200 μm for P0 and 500 μm for P6. (d) Violin plot for annotation only with P0 astroglial lineage sub-clusters. (e)-(h) Feature plot of BG-like cell specific markers (e, f) and proliferative markers (g, h) with P0 astroglial lineage cell sub-clusters. (i) Violin plot for annotation only with P6 astroglial lineage sub-clusters. (j)-(m) Feature plot of BG-like cell specific markers (j, k) and proliferative markers (l, m) with P6 astroglial lineage cell sub-clusters.

**Supplemental Figure 3.**
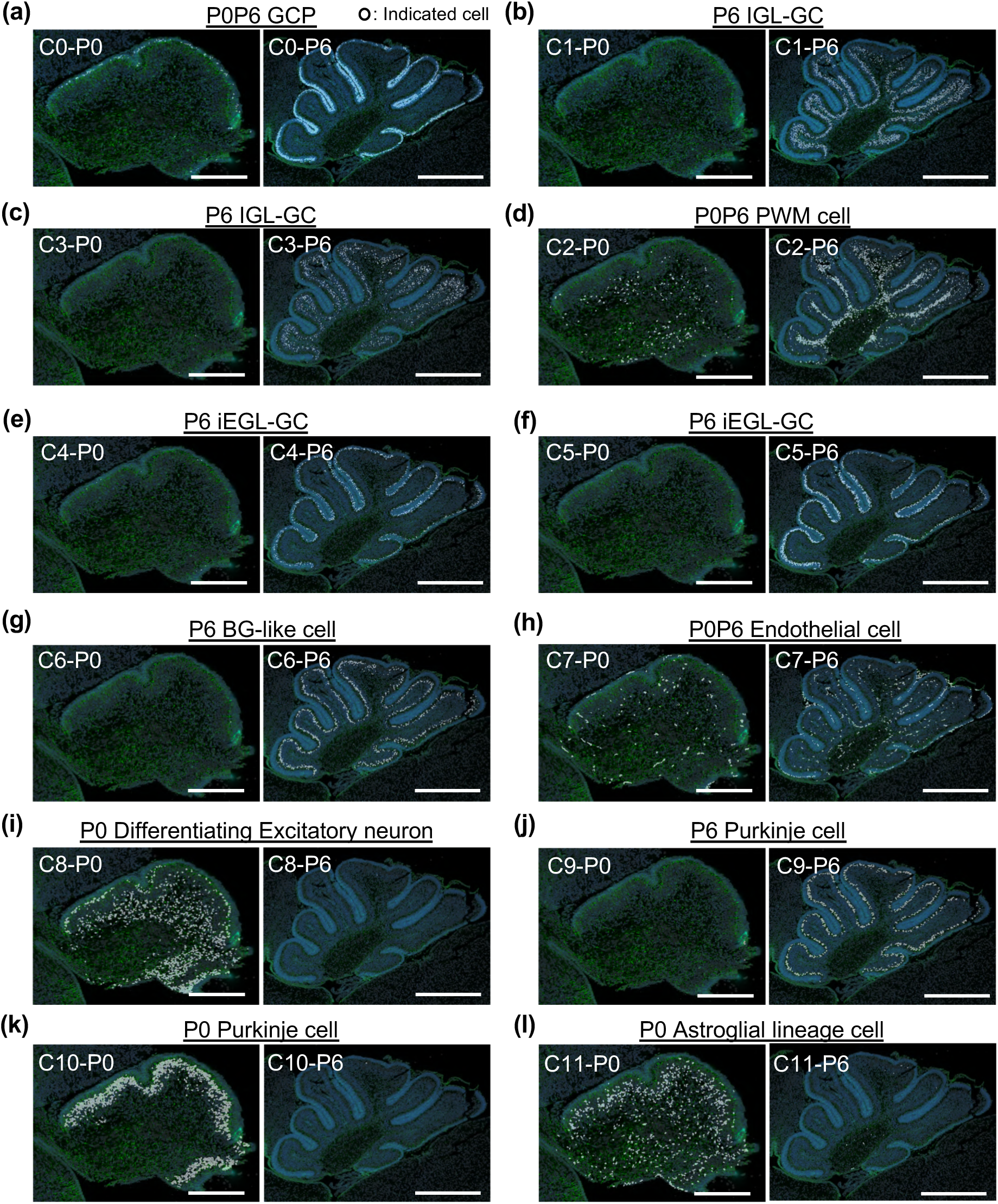
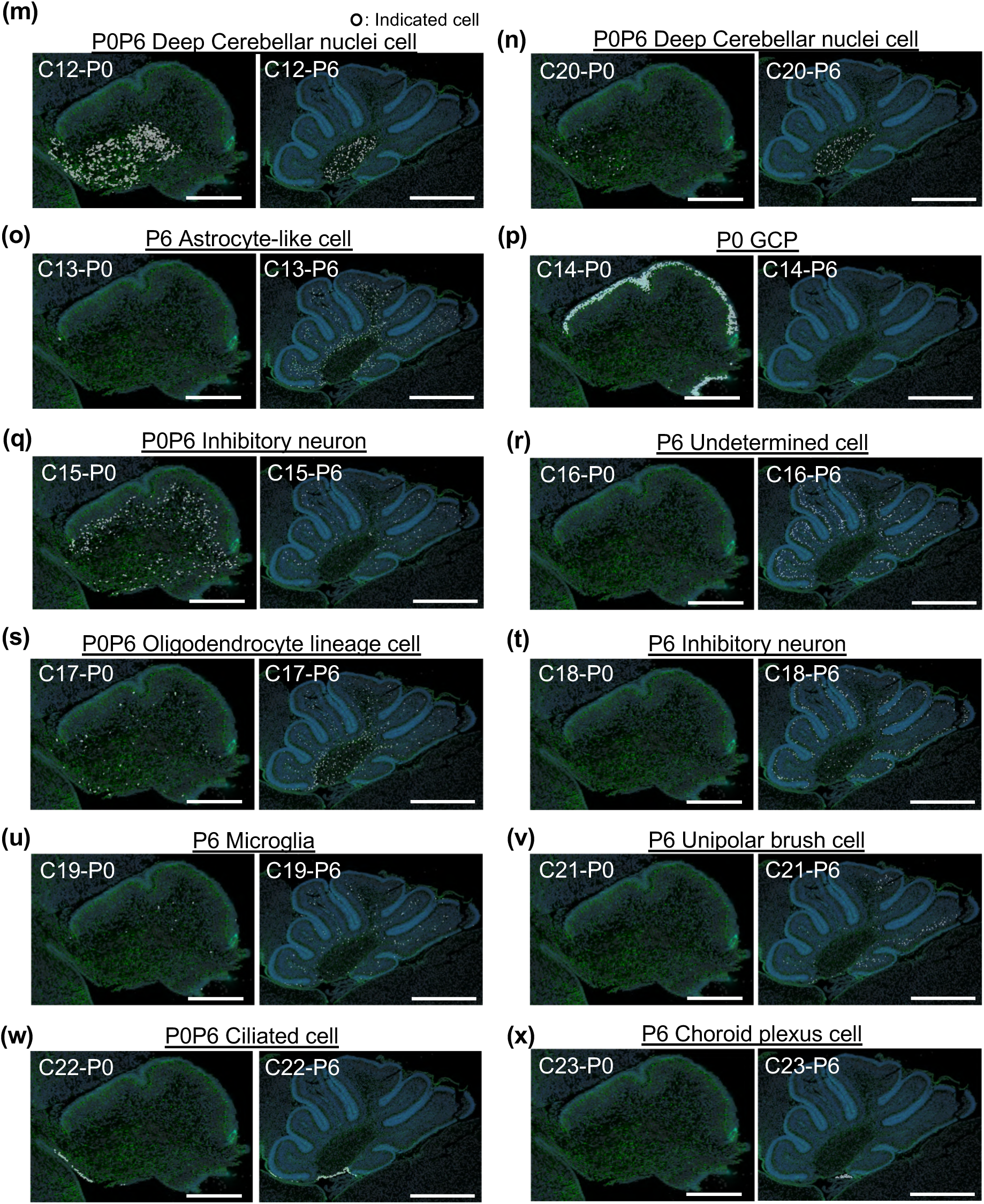
Spatial information of each 24 cluster. (a)-(x) Spatial localization information of each 24 cluster at each two stage which was used for annotation. Scale bars: 200 μm (left panels, P0) and 500 μm (right panels, P6).

